# LEVELNET to Visualise, Explore and Compare Protein-Protein Interaction Networks

**DOI:** 10.1101/2021.07.31.453756

**Authors:** Yasser Mohseni Behbahani, Paul Saighi, Flavia Corsi, Elodie Laine, Alessandra Carbone

## Abstract

Physical interactions between proteins are central to all biological processes. Yet, the current knowledge of who interacts with whom in the cell and in what manner relies on partial, noisy, and highly heterogeneous data. Thus, there is a need for methods comprehensively describing and organising such data. LEVELNET is a versatile and interactive tool for visualising, exploring and comparing protein-protein interaction (PPI) networks inferred from different types of evidence. LEVELNET helps to break down the complexity of PPI networks by representing them as multilayered graphs and by facilitating the direct comparison of their subnetworks toward biological interpretation. It focuses primarily on the protein chains whose 3D structures are available in the Protein Data Bank. We showcase some potential applications, such as investigating the structural evidence supporting PPIs associated to specific biological processes, assessing the co-localisation of interaction partners, comparing the PPI networks obtained through computational experiments versus homology transfer, and creating PPI benchmarks with desired properties.

**Availability**: LEVELNET is freely available to the community at http://www.lcqb.upmc.fr/levelnet/.

## 1 Introduction

Most of the cellular machinery and biological processes are fulfilled by biomolecular associations, particularly protein-protein interactions (PPIs). The versatile, adaptable, and specific nature of proteins gives them the ability to create networks that govern virtually all intra- and inter-cellular activities. Analyzing these networks enlightens the relative importance of proteins in different organisms and communities. It improves our understanding of physiopathological mechanisms and helps to decipher gene-disease-drug associations and find therapeutic treatments [29, 38, 35, 15, 42]. The ever increasing growth of protein sequential, structural, and functional information [44, 43, 4, 12, 17, 45] and of experimental evidence for PPIs [31, 30, 34, 32, 18] has stimulated the development of databases and web-interfaces for curating, inferring, and browsing PPI networks [6, 27, 28, 20, 8, 11, 19, 14, 23, 40]. These resources compile experimental and computational data and provide user-interactive visualization. A handful of them build upon the 3D structural evidence contained in the Protein Data Bank (PDB) [3], *e.g*. Interactome3D [27], PPI3D [8], and 3did [28]. Moreover, to cope with the biases, noise and uncertainty inherent to PPI-related data [13, 19], a lot of effort has been devoted to improve the reliability and interpretability of the inferred PPI networks, by integrating other types of information (*e.g*. cellular localisation), by organizing the PPIs based on their biological context, and by computing confidence scores. These characteristics are implemented in IID [19], STRING [41, 39, 40], and the human-focused base HIPPIE [2].

Here, we report on LEVELNET, a versatile computational framework designed to integrate and explore PPI networks coming from multiple sources of evidence. Starting from a set of protein chains whose 3D structures are available in the PDB, LEVELNET builds a grid of networks for each source (**Fig. 1**AB) representing different “views” on the interactions. It allows for clustering groups of similar proteins (nodes in the network) by exploiting global sequence identities between proteins and inferring interactions (edges in the network) through homology transfer or confidence scores (**Fig. 1**B). Networks coming from different sources can be integrated in an aggregated graph (**Fig. 1**CDG) where an interaction between two chains is represented as a multi-edge between two nodes, where the multiplicity comes from the different sources of evidence. Also, each edge is assigned a weight reflecting either a property from the source or the reliability of the evidence. This resulting information-rich framework helps to reason about interactions and to extract various biological information.

**Figure 1:**
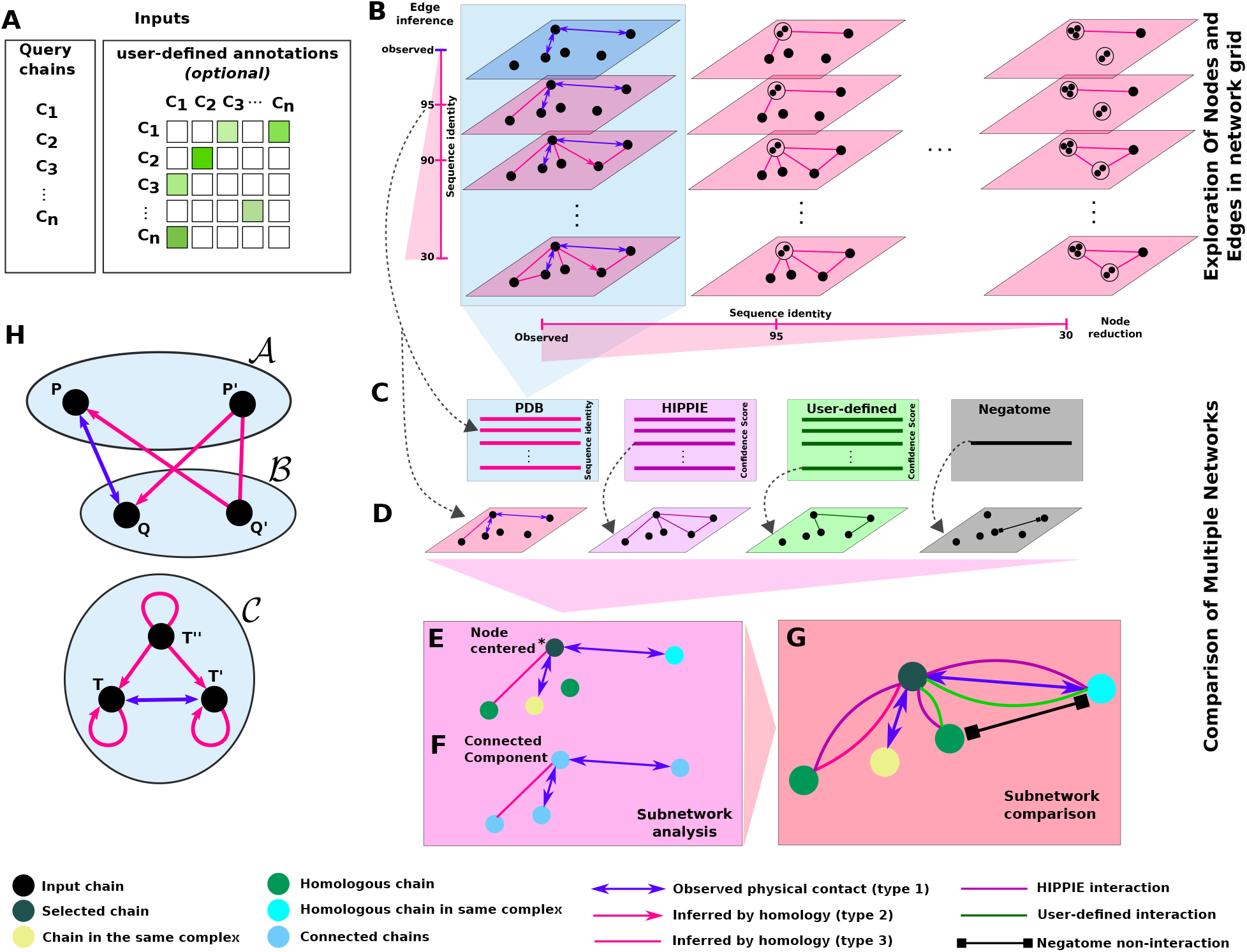
PPI network representation and analysis in LEVELNET. **A**. The input is a list of proteins (or protein chains) optionally accompanied by some pre-defined relationships (user-defined annotations). **B**. A grid of networks computed by LEVELNET from the PDB as source. The user has access to the layers by modulating the sequence identity on the “node reduction” and “edge inference” options and can visualize different networks of the grid. Similar grids are available for HIPPIE, Negatome, and user-defined sources (see Methods). **C**. Multi-layered networks from different sources share the same nodes (black dots) or super-nodes (sets of black dots) and correspond to a column of networks in the grid (see blue background in B). **D**. Selection and comparison of layers from different sources. **E**. Selection of a node-centered subnetwork in subnetwork analysis (the central node is shown by *). **F**. Selection of connected components in subnetwork analysis. **G**. Aggregated representation of the subnetwork in EF. **H**. Schematic representation of the inference of interactions from the PDB. 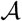, 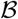, and 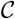 are clusters at some level of sequence identity, containing chains *P* and *P′*, *Q* and *Q′*, *T*, *T′* and *T″*, respectively. Chains *P* from cluster 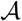 and *Q* from cluster 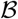 are in physical contact (blue edge). This interaction leads to inferring some interactions with and among their homologs (pink edges). When two chains from the same cluster are in direct contact, here *T* and *T′* from cluster 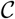, self-interactions are also induced by homology.

LEVELNET considers both interactions and non-interactions and represents them in different ways, thereby facilitating the identification of potential inconsistencies between different PPI sources. More specifically, it exploits inter-chain physical contacts extracted from the PDB, annotated interactions from HIPPIE and non-interactions from Negatome [36], and optionally user-defined interactions and/or non-interactions (**Fig. 1**C). It extends the set of direct physical interactions observed in the PDB among the input proteins by transferring knowledge from complex structures involving the same or very similar proteins. This choice is motivated by the observation that functional interactions are conserved across closely related homologs [25]. Moreover, works by us and others showed the biological pertinence and usefulness of accounting for homology-transferred interactions when evaluating protein-protein/DNA/RNA interface prediction methods [10, 47].

LEVELNET can be used to gain some visual interactive insight into a broad range of PPI-related questions, such as to what extent a signaling pathway or biological process has been structurally covered experimentally. Or where the interactions at play in a pathway take place in the cell and which proteins establish connections between the different cellular compartments. Or which of the PPIs predicted by an *ab initio* approach are supported by structural- and homology-based evidence. We illustrate these applications on the photosynthesis process, the Wnt signaling pathway, and a couple of established protein docking benchmarks. We show a few more usage cases, such as identifying cross-interactions among a set of complex structures or creating a set of interacting protein pairs devoid of such cross-interactions. Furthermore, to ease visualization, LEVELNET spatially arranges the nodes according to the topology of the network. On top of the grid-based visualisation, exploration and comparison of PPI networks, LEVELNET offers the possibility to retrieve detailed information, at the amino acid residue level, about the surface usage of a given protein in the PDB.

## 2 Materials and methods

### 2.1 PDB, HIPPIE and Negatome databases

PDB (June 2020 release) entries were downloaded from the FTP archive rsync.wwpdb.org::ftp/data/biounit. Entries with more than 100 chains or with a resolution lower than 5Å were discarded. Protein chains smaller than 20 residues or with more than 20% of unknown residues were also discarded. The HIPPIE database (v2.2) was downloaded from http://cbdm-01.zdv.uni-mainz.de/~mschaefer/hippie/download.php and Negatome database (v2.0) from http://mips.helmholtz-muenchen.de/proj/ppi/negatome/.

### 2.2 Pre-computed databases derived from the PDB

To infer interactions from the PDB, LEVELNET relies on two pre-computed databases. We describe here the memory efficient computational procedure we implemented to build these databases.

#### 2.2.1 Database of interfaces

We processed all biological assemblies (from X-ray crystallography and cryogenic electron microscopy) or NMR models from the PDB using the interface detection algorithm INTBuilder [9]. Two residues were considered as in contact if the distance between any of their atoms was smaller than 5Å. We call the resulting database of interfaces *interfaceDB*.

#### 2.2.2 Database of PPI networks

We pre-computed a grid of 6 × 6 = 36 PPI networks based on sequence identities between PDB chains (**Figure 1B**). We considered either all PDB chains individually (**Figure 1B**, see label *observed*) as *nodes*, or groups of chains defined at 5 levels of sequence identity, namely 30, 50, 70, 90 and 95% as *super-nodes*. To do this, we exploited the information contained in *interfaceDB* and the clusters of similar protein chains computed by MMSeqs2 [37] and available from the PDB. Formally, if

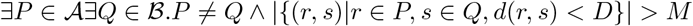

then the following edges will be inferred in LEVELNET (**Fig. 1**H):

- 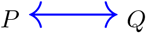 (type 1: double-arrow blue edge)
- for all 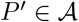 and 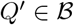.*P* ≠ *P′* and *Q* ≠ *Q′*
  – 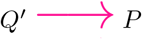 (type 2: pink directed edge)
  – 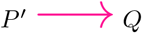 (type 2: pink directed edge)
  – 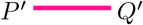 (type 3: pink undirected edge)

where 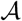 and 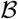 are two clusters of chains in a certain percentage of sequence identity (**Fig. 1**H), *d*(*r, s*) defines the distance between two residues *r* and *s*, *D* is the distance threshold set to 5Å by default, *M* is the minimum number of interface residues set to 5 for both proteins by default. Note that this formalism is also suitable for inferring interactions within a given cluster (*i.e*. when 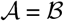, see **Fig. 1**H, bottom panel).

In practice, our algorithm first loops over all pairs of chains in all PDB complex structures to verify, for each pair, the existence of a contact in *interfaceDB* and infer a *chain-chain* interaction. Such an interaction is of type 1. Then, it loops only over the identified directly interacting chain pairs (*P, Q*) and determine sets of *chain-cluster* and *cluster-cluster* interactions, for each level of sequence identity considered:

1. **foreach** pair (*P, Q*) **do**
2. 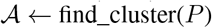
3. 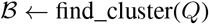
4. 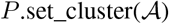
5. 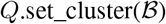
6. 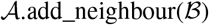
7. 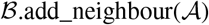
8. 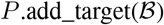
9. 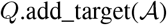
10. **end**

On lines 2 and 3, the function *find*_*cluster* determines to which clusters chains *P* and *Q* belong at a certain percentage of sequence identity. Lines 4 and 5 set the cluster identifiers as the chains’ attributes. On lines 6 and 7, the function *add*_*neighbour* sets a cluster-cluster interaction. This interaction implies that all members of 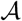 will be linked to all members of 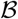 by interactions that are at least of type 3. On lines 8 and 9, the function *add*_*target* sets two chain-cluster interactions. This operation implies that *P* (resp. *Q*) will be linked to all chains from 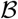 (resp. 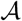) by interactions of at least type 2. We call the resulting database of chain-chain, chain-cluster and cluster-cluster interactions *PDBinteractionDB*.

### 2.3 Description of LEVELNET functionalities

#### 2.3.1 Operational modes and inputs

LEVELNET operates in two modes. In the **query-versus-query** mode, it allows interactively exploring and comparing the interactions among a set of input proteins or proteins chains, designated by their PDB identifiers. The users can additionally provide an input matrix specifying the existence or non-existence and the strength or confidence of some relationships between them (**Fig. 1**A). The matrix should be given as a list of triplets: protein1, protein2, and associated score of interaction (i.e., a value between 0 and 1; if a score is missing, the value 1 is taken by default). In the **query-versus-all** mode, LEVELNET allows retrieving residue-level information about the interactions established between a set of input protein chains, designated by their PDB identifiers, or their homologs, and all other protein chains in the PDB.

#### 2.3.2 Query-versus-query mode

In the query-versus-query mode, given a set of proteins, LEVELNET assigns a grid of networks to each source of evidence. In case of the PDB source, this grid comprises 36 PPI networks as described above but restricted to the input proteins. In case of HIPPIE and Negatome databases and the user-defined input matrix, LEVELNET creates a grid of 6 × *K* networks based on five levels of sequence identity for the nodes (30, 50, 70, 90 and 95%) and the “observed” one, and on *K* running on all confidence score values for the edges.

We define six types of edges:

- 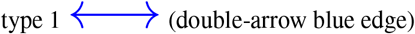, observed interaction, *i.e*. the two chains are in physical contact in a known complex PDB structure,
- 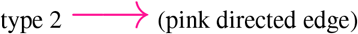, interaction inferred by homology where one of the chains is in physical contact with a homolog of the other chain in a known complex PDB structure,
- 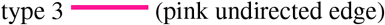, interaction inferred by homology where some homologs of the two chains are in physical contact in a known complex PDB structure,
- 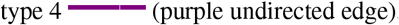, HIPPIE annotated interaction,
- 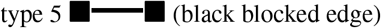, NEGATOME annotated non-interaction,
- 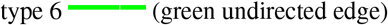, user-defined interaction.

Given an input set of protein chains, LEVELNET will interrogate *PDBinteractionDB* to create type 1, 2 and 3 edges. For each chain pair in the set, if there is a chain-chain interaction, LEVELNET creates a type 1 edge. If not, then if there is a chain-cluster interaction it creates a type 2 edge, otherwise if there is a cluster-cluster interaction, it creates a type 3 edge. LEVELNET will also interrogate HIPPIE and Negatome to infer type 4 and 5 edges. Finally, if the user specified some relationships between the input chains, LEVELNET will integrate them in the output multi-layered network as type 6 edges. Moreover, LEVELNET annotates some of the edges with weights representing either the level of sequence identity at which the corresponding interaction was inferred (type 2 or 3), a confidence level (type 4), or any type of user-defined annotations (type 6). Each network layer comprises the set of edges inferred from a single source and with a weight higher than a certain threshold (**Fig. 1**B).

##### Network exploration

The users can navigate from one layer to another within the grid associated with each source and also across the grids. Within a grid, they can modulate both the number of nodes and the number of edges. Upon relaxing the node sequence identity threshold, the nodes representing similar chains will be progressively merged into *super-nodes* and thus the network will simplify. Upon relaxing the edge sequence identity or confidence threshold, new edges will appear and thus the network will become denser.

##### Subnetwork comparison

LEVELNET allows for comparing several layers, focusing on a selected subset of input proteins (**Fig. 1C**). The selection can be:

- *node-centered*, upon clicking on a chosen node. LEVELNET then highlights its homologs in green and the chains belonging to the same complex in yellow (**Fig. 1**D). This functionality helps, for instance, to detect homo-oligomers.
- *at the level of a connected component*, to focus on a signaling pathway for instance (**Fig. 1**E).

Once the users have selected a subset of nodes, they can create a multi-edge graph by superimposing several layers and directly compare the corresponding interactions (**Fig. 1**G). This analysis also allows the users to discover inconsistencies among various resources of PPI.

#### 2.3.3 Query-versus-all mode

Beyond allowing for the identification, visualisation and comparison of PPIs, LEVELNET provides the users with a residue-level description of protein interaction surfaces. In this operational mode, the web server outputs the ensemble of interacting patches, each query chain and its homologs (at a certain level of sequence identity) together with all chains in the PDB that are physical partners in some complex. The interacting patches corresponding to the physical interactions are mapped onto the input query chain and merged to provide a label, either interacting or non-interacting, to each surface residue of the query protein.

#### 2.3.4 Implementation details

To generate the PDB interaction database, we developed a pipeline in Python that processes PDB protein complexes, their interfaces, and chain clusters by sequence identity. We created an interactive environment based on recent advances in web development including HTML5 and related technologies such as D3, JavaScript, Vue, and SVG. HTML5, CSS3 and Vue are used for the front-end and provide a stylish and user-friendly interactive interface. Data visualization is performed using D3 [5] and SVG. Codes developed in JavaScript and Python process databases and user queries. The whole pipeline is optimized to respond to queries rapidly.

### 2.4 Benchmark databases used in the applications

We used three benchmark databases to showcase the applications of LEVELNET. These databases are the Docking Benchmark ZDock version 5.5 (ZDockv5) [46] and version 2.0 (ZDockv2) [26] and DockGround (DG4) [21]. We used all the single-chain proteins of these databases. For ZDockv2, we chose a subset of 88 single-chain proteins (out of 168 proteins) for our evaluation purposes (see Results) and called it ZDockv2_s88.

## 3 Results and discussion

### 3.1 Exploring and discovering the interactions underlying photosynthesis

As a first case study, we considered photosynthesis in the green alga *Chlamydomonas reinhardtii*. The structural information available for this process is very partial, and only a few protein complexes are resolved in the PDB (see Supplementary Materials and Methods). LEVELNET computed a PPI network comprising 1 430 pairwise interactions between 108 structurally resolved protein chains (**Fig. 2**). The network is characterized by three large connected components that are very easy to delineate thanks to LEVELNET’s ability to spatially group inter-connected nodes (**Fig. 2**A, see encircled regions). The connected components correspond to the photosystem light harvesting complexes (PLHC) of types I and II, respectively, and the Ribulose-1,5-bisphosphate carboxylase-oxygenase (RuBiSCo) complex. Upon merging the nodes sharing more than 95% sequence identity, the RuBiSCo component reduces to only two super-nodes corresponding to the small (S, in red) and large (L, in blue) subunits of this key enzyme of the Calvin-Benson-Bassham cycle (**Fig. 2**B). LEVELNET’s highlighting of the nodes belonging to the same PDB complex allows understanding who is connected to whom in the Cytohrome b6f enzyme (**Fig. 2**A, see nodes in green and yellow, and (**Fig. 2**C). Overlaying the information coming from the PDB and the Negatome allows spotting inconsistencies between these databases (**Fig. 2**C).

**Figure 2:**
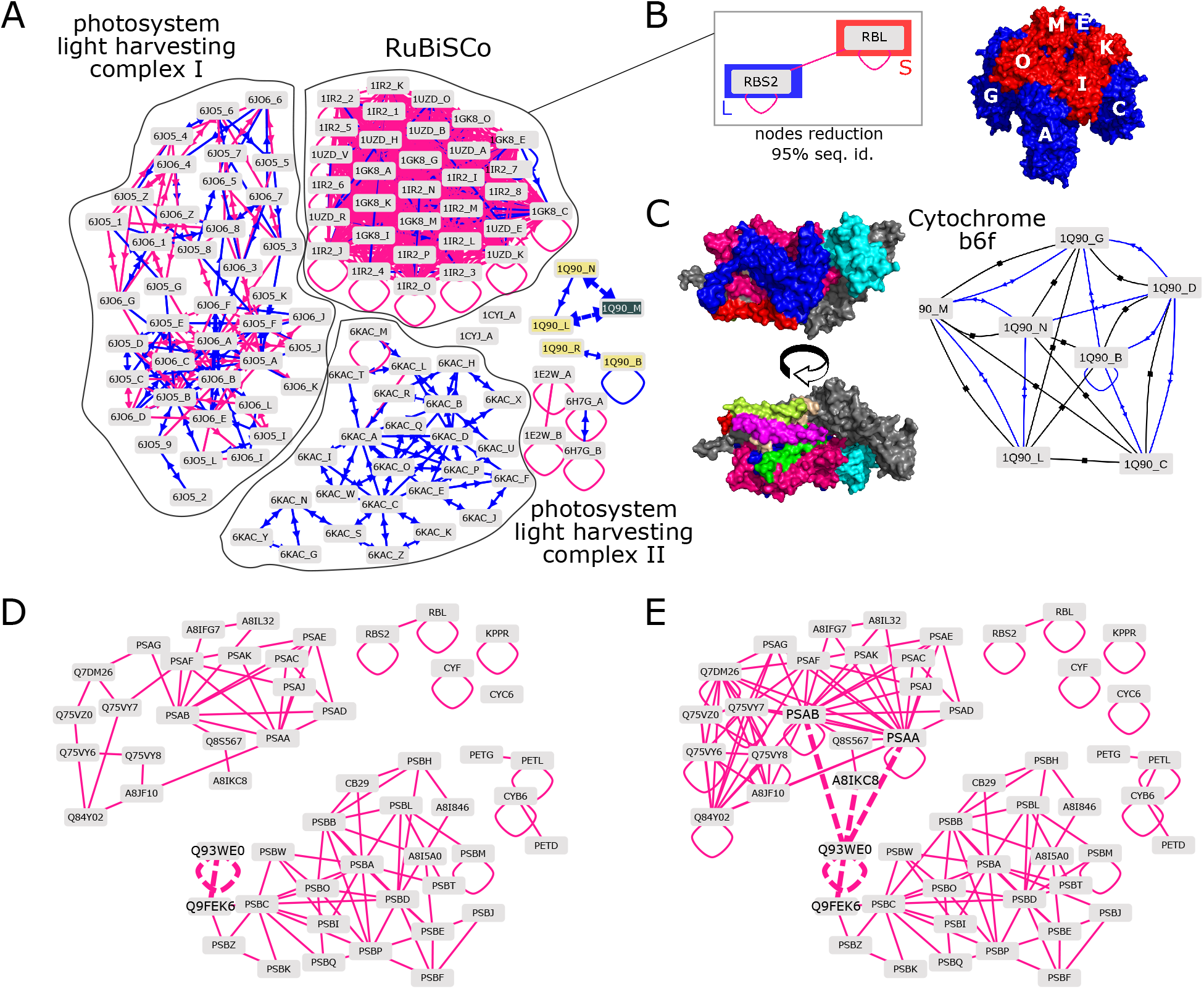
Structurally characterized PPI network for photosynthesis in *Chlamydomonas reinhardtii*. **A**. Connected components identified in the PPI network computed using the PDB as the source database. Each node represents a protein chain. The blue edges represent observed interactions while the pink ones represent homology-transferred interactions at more than 95% sequence identity. The connected components correspond to the RuBiSCo complex, and the photosystem light harvesting complexes of types I and II. This network also includes other protein chains forming small subnetworks. **B**. Node reduction on RuBiSCo: merging the nodes with more than 95% sequence identity results in two super-nodes corresponding to the small and large subunits of the RuBiSCo complex 3D structure, in red and blue respectively. **C**. Subnetwork comparison of two layers, corresponding to the interactions observed in the PDB (blue) and the non-interactions from the Negatome (in black) for the Cytochrome b6f enzyme. Two views of the Cytochrome 3D structure. **D-E**. Interaction networks after merging nodes sharing more than 70% sequence identity: edge inference by 95% (**D**) and 30% (**E**) sequence identity.

Setting the sequence identity threshold at 70% for node reduction and 95% for edge inference from the PDB leads to a network comprised of 96 pairwise interactions between 53 nodes (**Fig. 2**D). Relaxing the threshold to 30% for edge inference densifies the PLHC-I subnetwok, and reveals a few new self-interactions and also cross-interaction between PLHC-I and PLHC-II (**Fig. 2**E). In particular, three interactions are inferred between the LHCII antenna (comprised of LHCII-1.3, LHCII-3 and LHCII-4 and represented by the node Q93WE0) from the PLHC-II subnetwork and three proteins from the PLHC-I subnetwork, namely the P700 chlorophyll a apoproteins PSAA and PSAB and the Chlorophyll a-b binding protein LHCA2. This inference is consistent with the fact that LHCII acts as an antenna for both photosystems I and II [7, 16] and in perfect agreement with a series of recent PDB structures of the supercomplex PSI-LHCI-LHCII from *Chlamydomonas reinhardtii*, released after June 2020 and thus not included in the present analysis [33].

### 3.2 Localizing PPIs in the Wnt signaling pathway

Next, we focused on the canonical Wnt signaling pathway that regulates gene transcription by passing signals from the cell surface receptors to the nucleus [1]. We used LEVELNET to visualise the structurally determined PPIs involved in this pathway in the context of their subcellular localisation. As input, we considered the list of 85 PDB entries reported in [1] and defined a custom adjacency matrix linking the protein chains localized in the same cellular compartments. LEVELNET built a network comprising 235 nodes linked by 2734 edges representing binary interactions observed in the PDB between the input chains or their homologs at more than 30% sequence identity. Based on the input custom matrix, LEVELNET automatically determined some spatial arrangement for the nodes (**Fig. S1**). In the resulting network, one can clearly identify groups of proteins sharing the same location and functional role, and distinguish the interactions taking place within each group from those taking place between the groups. Upon merging the nodes sharing more than 30% sequence identity, the size of the network reduces dramatically down to less than 50 nodes. This non-redundant version allows reasoning at the protein level, toward identifying the parts of the pathway that are described by structural information and the parts where such information is missing (**Fig. S1**, compare the inset with Figure 1 from [1]). Several complexes in the membrane have been described, for instance between Dickkopfs (DKKs), Kremen (KREM1) and LRP6, between R-spondins (RSPOs) and LGR family receptors, and between Wnt proteins (WNTs) and Frizzled receptors (FZDs) (**Fig. S1**, grey panel in inset, from left to right). By contrast, no structural information is available for the complex between Wnt and its negative regulators Wnt inhibitory factor and secreted-Frizzled related proteins (**Fig. S1**, red panel in the inset, see WIF1 and SFRP3 with self-edges only). *β*-catenin (CTNB1) plays a central role in the pathway, at the interface between the nucleus (blue panel) and the cytoplasm (green panel). In particular, in the cytoplasm (green panel), its interactions with axin (AXN) and Adenomatous Polyposis Coli (APC) contributing to the formation of the degradosome were structurally characterized. So were its interactions with the scaffold protein BCL9/legless, Groucho/TLE, and *β*-catenin binding proteins/domains (CNBP1, TCF3-CBD), taking place within the Wnt enhanceosome located in the nucleus (blue panel).

### 3.3 Comparing predicted versus homology-transferred PPIs

We used LEVELNET to compare the PPIs predicted by a complete cross-docking experiment [24] with the structurally characterized PPIs available in the PDB (**Fig. 3** and **S5**). We considered an ensemble of about 4 000 putative protein pairs coming from the ZDockv2_s88 (see Materials and Methods). The cross-docking experiment yielded high interaction strengths for 151 pairs (*NII* >= 0.7, see green curve in **Fig. 3**A). While only 12% of those are supported by the complex structures annotated in the benchmark (*Observed*, dark blue curve), this proportion increases up to 20% when considering all the complexes in the PDB formed between the chains of the benchmark set or their close homologs (at >90% sequence identity, petrol curve). Beyond providing global statistics, LEVELNET facilitates such comparison at the level of specific subnetworks. As an example, we consider a PPI subnetwork centred on the GTP-binding nuclear protein RAN (**Fig. 3**B). The information LEVELNET gathered from the PDB describes interactions between RAN (1A2K:C, and also 1IBR:A, 1K5D:A and 1I2M:A) and 5 partners, namely the nuclear transport factor 2 (1A2K:A and 1A2K:B), the Ran-specific GTPase-activating protein (1K5D:B), the Ran-binding protein 1 (1K5D:C), the Importin beta-1 subunit (1IBR:B) and the regulator of chromosome condensation RCC1 (1I2M:B) (**Fig. 3**B, see the blue and pink dotted edges between 1A2K:C and the grey or yellow nodes). Three of these interactions are correctly predicted by the cross-docking (green dotted edges). Moreover, both the PDB and the cross-docking experiment supports the formation of RAN homodimers (see the dotted edges between 1A2K:C and the green nodes). As expected, the docking calculations are sensitive to conformational changes, as illustrated by the interaction between RAN and the Importin beta-1 subunit: while the interaction is predicted when docking 1IBR:A against 1IBR:B, it is not predicted when docking 1A2K:C against 1IBR:B. The structure 1A2K:C differs from 1IBR:A by a completely different orientation for a 10-amino acid long loop and an extra C-terminal helix. Finally, one of the predicted interactions is in conflict with the Negatome layer (black edge, chains BC of 1K5D).

**Figure 3:**
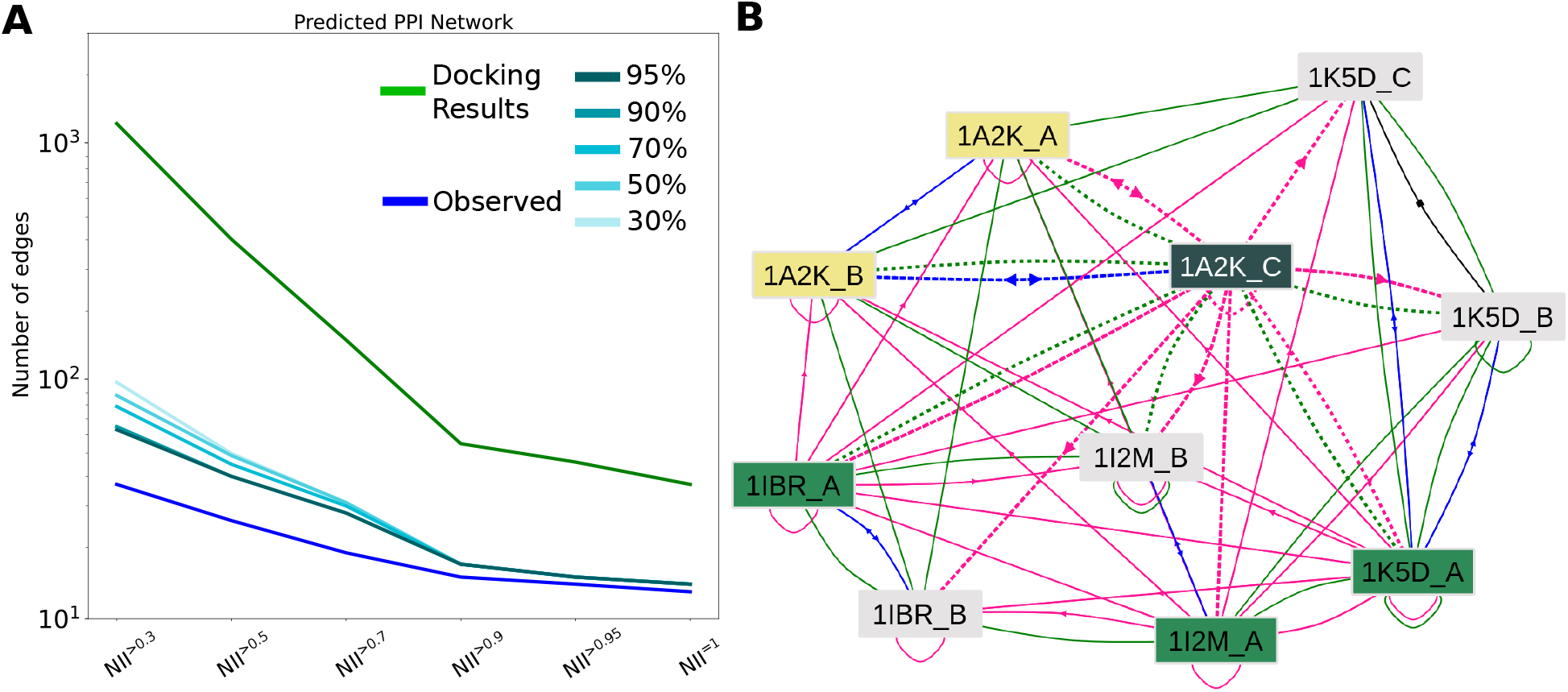
Assessments of PPI predictions. **A**. Number of PPIs predicted for a subset of 88 single-chain proteins from ZDockv2 through complete cross-docking simulations [24] and supported by experimental structural data. The normalized Interaction Index (NII) is a confidence score assigned to each putative protein pair. Note that when a protein contains several chains, all pairs of interacting chains are assigned a score of 1. We gave these scores as an input custom matrix to LEVELNET. The y axes are in logarithmic scale. **B**. Subnetwork for the GTP-binding nuclear protein RAN and its partners. The chains corresponding to RAN are 1A2K:C, 1IBR:A, 1K5D:A and 1I2M:A. Upon clicking on one of them, here 1A2K:C, LEVELNET interactive interface highlights the other ones, identified as homologs, in green. The other chains belonging to the same PDB complex are highlighted in yellow. The observed and homology-transferred (at >95% sequence identity) interactions are represented by the blue and pink edges, respectively. The green ones represent interactions predicted by cross-docking (taken from [22]. The interactions involving the selected chain (1A2K:C) are highlighted with dotted lines.

### 3.4 Decrypting and customising benchmarks

LEVELNET can be used to complement the information provided in a protein benchmark set. For instance, while each protein from the ZDockv5 benchmark [46] has only one annotated cognate partner, LEVELNET identifies up to 200 partners by homology transfer, at >70% sequence identity, from the PDB (Figures 4 and S2AC). The nodes with the highest degrees correspond to antigen-binding fragments (FAB) (**Fig. 4**A). This result reflects the very high sequence identity shared by FABs recognizing different antigens. Indeed, only a few highly variable residues on antigen-binding site forming the paratope are responsible for the specific recognition of the antigen (**Fig. 4**A, red sticks). In such cases, transferring interactions by homology may not be valid, even at 95% sequence identity. Leaving out this functional class, we still observe many homology-transferred interactions, with up to 40 partners for one single protein (**Fig. S2**A). The biggest connected components comprise several hundreds of proteins (Figures 4B and S2CD). A similar trend can be observed for the DockGround (DG4) (**Fig. S2**BD). This type of analyses helps to get a broader view of a set of proteins than the annotations at hand permit, and also emphasizes the complexity underlying the behaviour of a protein within a community.

**Figure 4:**
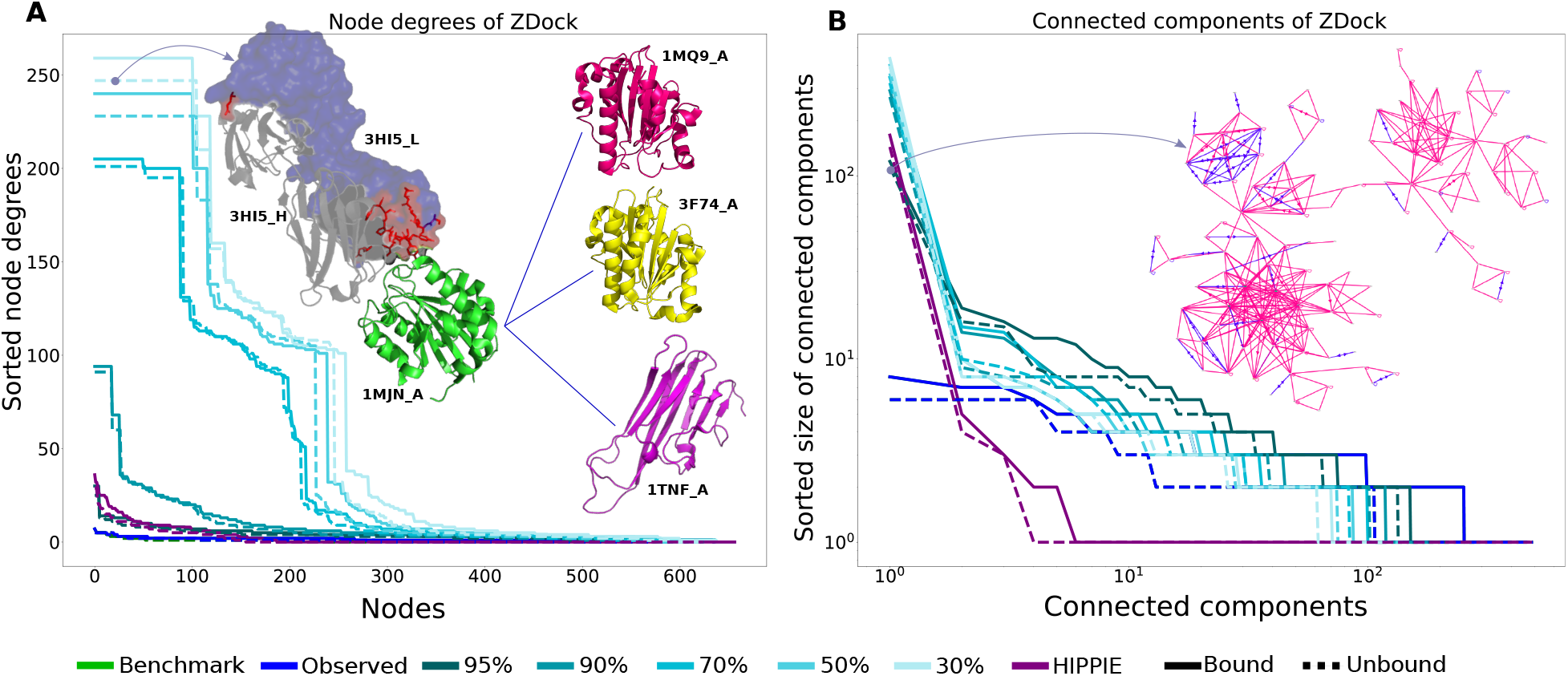
Node degrees and size of connected components in LEVELNET for ZDockv5. For each layer, the node degrees (**A**) and the sizes of connected components (**B**) are sorted from the largest to the smallest. **A**. The inset of the plot shows the human FAB structure of 3HI5, characterized by the highest node degree in the dataset. Residues with very high variations among the homologs of this FAB light chain are distinguished by red sticks. The majority of these residues belong to the paratope part of the light chain. By homology transfer at 95%, this FAB light chain has the potential to interact with other FAB heavy chains or antigens in the dataset, including 1MQ9, 3F74, and the Lymphokines trimer 1TNF. **B**. The inset of the plot shows the topology of the largest connected component, obtained at more than 95% similarity, including the FAB structure of 3HI5.

Beyond characterizing the properties of existing benchmark sets, LEVELNET can be effectively used to create new benchmarks with desired properties. To showcase this functionality, we selected a set of 500 high-quality hetero-trimeric structures from the PDB (see Methods) and gave it as input to LEVELNET. We focused on the network layer defined from the PDB at 70% sequence identity. It comprises 1206 chains and is organized into 124 connected components. Upon redundancy reduction, the network resumed to 112 connected components, among which 45 were made of only 3 nodes (represented by 3 master chains). This analysis shows that it is straightforward to compile new benchmarks with LEVELNET for assessing a specific PPI-related task, like predicting how 3 proteins assemble together. Our procedure guarantees that no cross-interaction in the set exists (based on the available structural information) and that the proteins differ by more than 30% from each other. The full list of chains from the set is given in the Supplementary Material. The PPI network of this benchmark is shown in **Supplementary Fig. S6**.

### 3.5 Interacting patches on the Glucocorticoid ligand-binding domain

We applied Query-versus-All functionality to the glucocorticoid receptor ligand-binding domain (Figures 5A). Starting from the input query chain 1NHZ:A, LEVELNET retrieved 61 interacting patches at 90% sequence identity. While the initial PDB complex structure 1NHZ displays an interacting patch of 33 residues (out of 239 in total), LEVELNET identified four times more (130) interacting residues by considering the whole PDB. This dramatic increase reflects the multiple binding modes with which this protein can self-assemble (Figures 5B).

**Figure 5:**
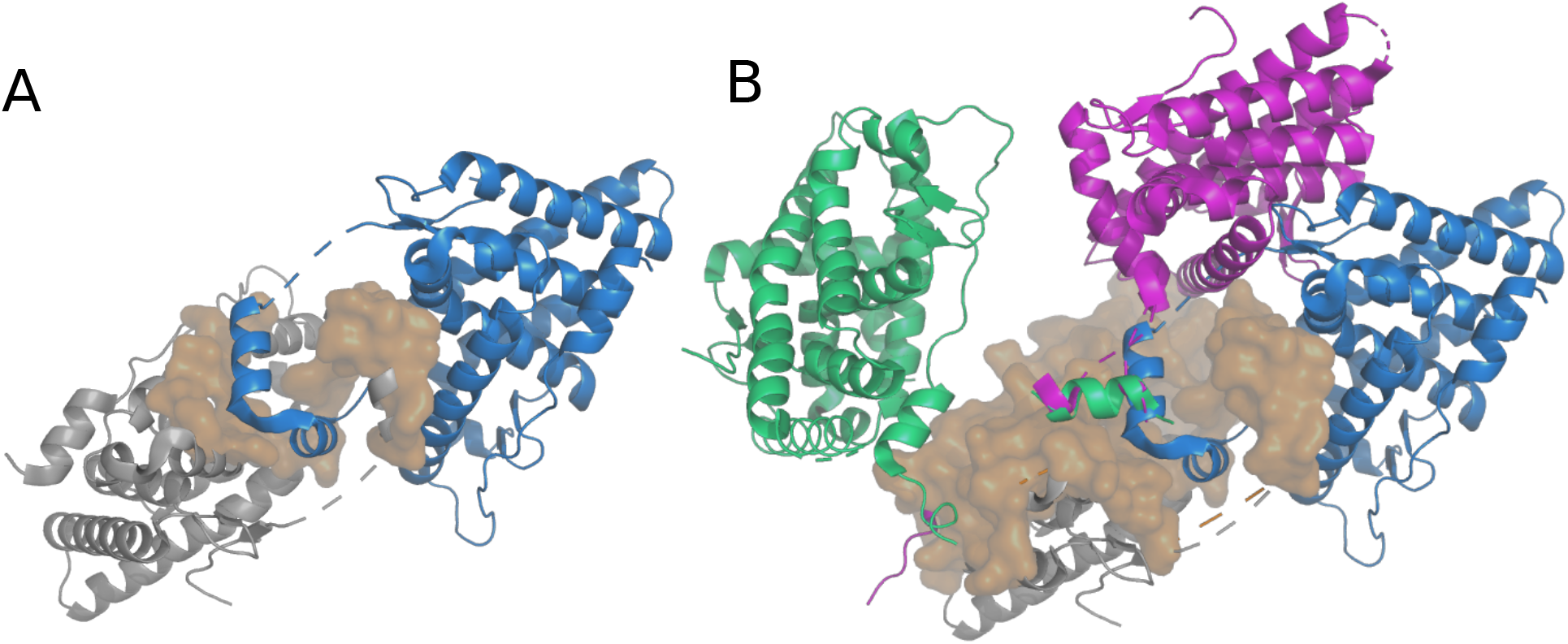
Interacting surfaces of the glucocorticoid receptor ligand-binding domain. The query protein chain (PDB id: 1NHZ:A) is displayed as a grey cartoon. **A**. Homodimer corresponding to the entry 1NHZ (second biological assembly). The interacting surface is highlighted in surface and the second copy of the protein is in marine cartoons. **B**. Different binding modes for the protein self-assembly. The highlighted surface covers the residues engaged in at least one interaction involving the query chain or a homolog at 90% sequence identity in the PDB. The blue, magenta and green cartoons represent interacting copies of the protein in the PDB complexes 1NHZ, 3E7C, and 4LSJ respectively.

### 3.6 Comparison with other PPI resources

LEVELNET allows the interactive and in-depth exploration of the interactions and sequence similarities shared among a set of proteins of interest. Compared to other web services dedicated to PPIs, it features a unique combination of structural information, homology transfer and annotations for interactions and non-interactions (Table 1). It offers the possibility to compare this information with user-defined annotations that can represent predicted interactions, co-localisation or any other type of relationship. It permits to dynamically visualise the evolution of a PPI network upon changing the edge confidence level. Compared to Interactome3D, LEVELNET has the advantage of providing PPI confidence scores and to handle homology transferred interactions. Compared to PPI3D, it has the advantage of featuring an interactive visualisation of PPI networks. Moreover, it processes user queries very rapidly by pre-building the interaction and physical contact databases. PPI3D has a longer response time due to PSI-BLAST queries performed for inferring interactions by homology.

**Table 1:**
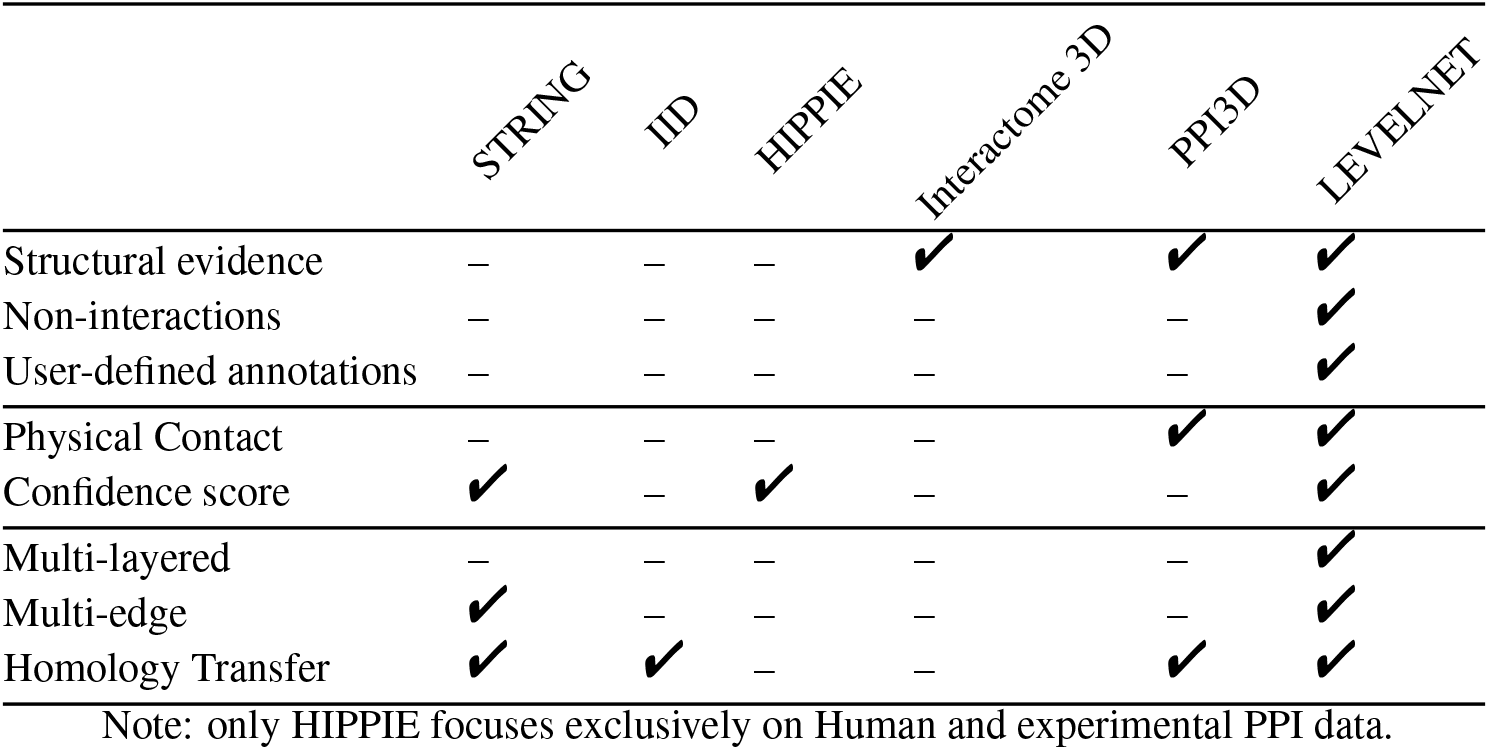
Comparison between PPI resources

## 4 Conclusion

LEVELNET is a valuable asset for the community to explore protein interactions. It is useful for the biologists interested in the physical contacts of a particular protein or a set of proteins as well as for those who develop and assess computational predictive approaches for interface, partner and complex predictions. It provides a convenient mean to account for different types of relationships between proteins, *e.g*. functional annotations, cell co-localization, spatiotemporal proteomics, or co-occurrences in publications, and investigate how the latter are correlated to physical interactions. We performed all the analyses reported here based on the June 2020 release of the PDB. Nevertheless, we have been updating LEVELNET based on more recent releases and will continue to maintain the pre-computed databases up-to-date. Future developments will concern the integration of other databases, be they computational or experimental. In particular, we will upgrade LEVELNET with information coming from reliable predictions of protein complexes using AlphaFold, RosettaFold or methods inspired by the latter. We expect the body of these predictions to massively increase in the coming years. Another direction will be to allow for merging and selecting nodes based on annotations from Uniprot, SCOP or CATH. LEVELNET could be improved by implementing the possibility to deal with multi-chain proteins and protein-nucleic acid interactions. Finally, we plan to increase the richness of the information provided by LEVELNET, *e.g*. by describing at the residue-level the binding sites and the binding-associated conformational changes, and giving access to properties such as binding affinities and the effect of mutations over the network.

## 5 Availability

LEVELNET is freely available to the community at http://www.lcqb.upmc.fr/levelnet/.

All datasets used and generated in this study are made available to the community at http://www.lcqb.upmc.fr/levelnet/#/tutorial.

## Acknowledgements

We thank Francesco Oteri for the technical support and Julien Henri for helping us with the analysis of photosynthesis process in the *Chlamydomonas reinhardtii*.

## SUPPLEMENTARY INFORMATION

### Supplementary Materials and Methods

#### Details about the pre-processing of the input queries

##### Protein chains involved in the photosynthesis from the green alga *Chlamydomonas reinhardtii*

To obtain all the proteins from *Chlamydomonas reinhardtii* involved in photosynthesis, we first performed a search in UniProt with the following criteria: “Chlamydomonas Reinhardtii [3055]” as organism and “photosynthesis [15979]” as Gene Ontology (GO) annotation. Next, we chose one PDB entry for each UniProt code of this list. We took the PDB entry with the maximum coverage of the sequence in the cases where multiple PDB codes are associated to a UniProt code. Finally, we queried the resulted set against LEVELNET database.

##### Benchmark creation in Section 3.4

To create a new benchmark set, we first performed an advanced search in the PDB. The criteria for this search were the following: protein as Polymer Entity Type, Hetero 3-mer as Oligomeric State, Polymer Entity Sequence Length between 100 and 300, Refinement Resolution better than 2.5 Å. Moreover, all complexes related to antibody-antigen (e.g. FAB) were excluded.

#### Discovery of self- and cross-interactions

The variation of the topology of the networks with respect to the interactions between each layer and corresponding benchmark network is shown in **Supplementary Fig. S3**. Self- and cross-interactions are considered separately and the difference is calculated as the percentage of the maximum possible connections: *n* and 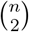 (fully connected graph), respectively. The only difference between physical contact and benchmark networks is the self-interactions. These self-interactions are caused by the identical copies of the same chain found in a biological assembly that are in contact. For both benchmarks, more than half of the nodes can interact with themselves at sequence identity above 95%. In case of cross-interactions, ZDockv5 with AA and AS included shows higher differences for both unbound and bound annotations compared to other benchmarks. As a further comparison we studied the pairwise differences in the number of interactions for layers of LEVELNET before and after redundancy reduction (**Supplementary Fig. S4**).

**Supplementary Fig. S7** displays a snapshot of the observed and homology transfered interactions with > 95% sequence identity of single-chain proteins of the ZDockv5 in the bound annotation. The users can highlight and isolate the nodes of the connected protein chains by searching their codes (**Supplementary Fig. S7**D) (the blue connected component subnetwork) as well as follow the trace of homologous chains by clicking on each chain (the node-centered subnetwork) (**Supplementary Fig. S7**A). The essential information of each protein chain will be displayed in a pop-up window by hovering mouse on its node. For a better visualisation the users can control the attractive/repulsive force between the nodes to discriminate clusters of densely connected proteins and also change the edge length, width, strength and node sizes to regulate the dynamic and floating property of the network (**Supplementary Fig. S7**C). The users can visualize another layer of LEVELNET by changing the source, modulating the confidence score, or choosing the non-redundant representation (**Supplementary Fig. S7**B). LEVELNET does not require any plugins; and is compatible with all web browsers. User can download, export, or share the PPI network. LEVELNET is accessible on smartphones and tablets to facilitate the collaboration between researchers.

**Figure S1:**
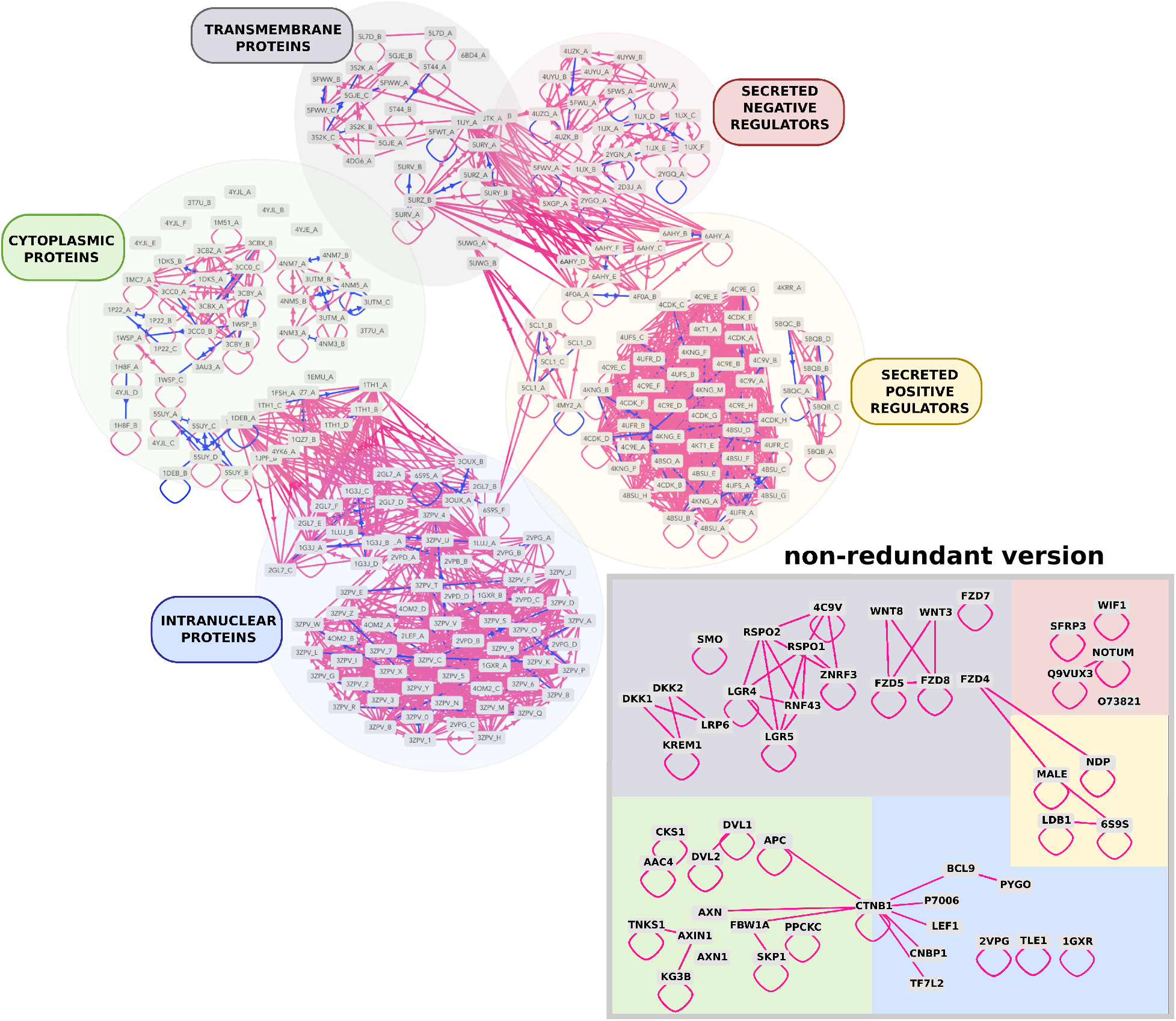
Network inferred for the canonical Wnt signaling pathway. The input PDB codes were taken from [1], and we show the layer corresponding to the interactions inferred from the PDB at 30% sequence identity. The co-localization annotations of the proteins taken from [1] were given as part of the input, and LEVELNET automatically determined the spatial arrangement of the nodes based on these annotations. The groups corresponding to the different protein cellular locations and roles are labelled. The colors of the circles around the labels are also used in the inset.

**Figure S2:**
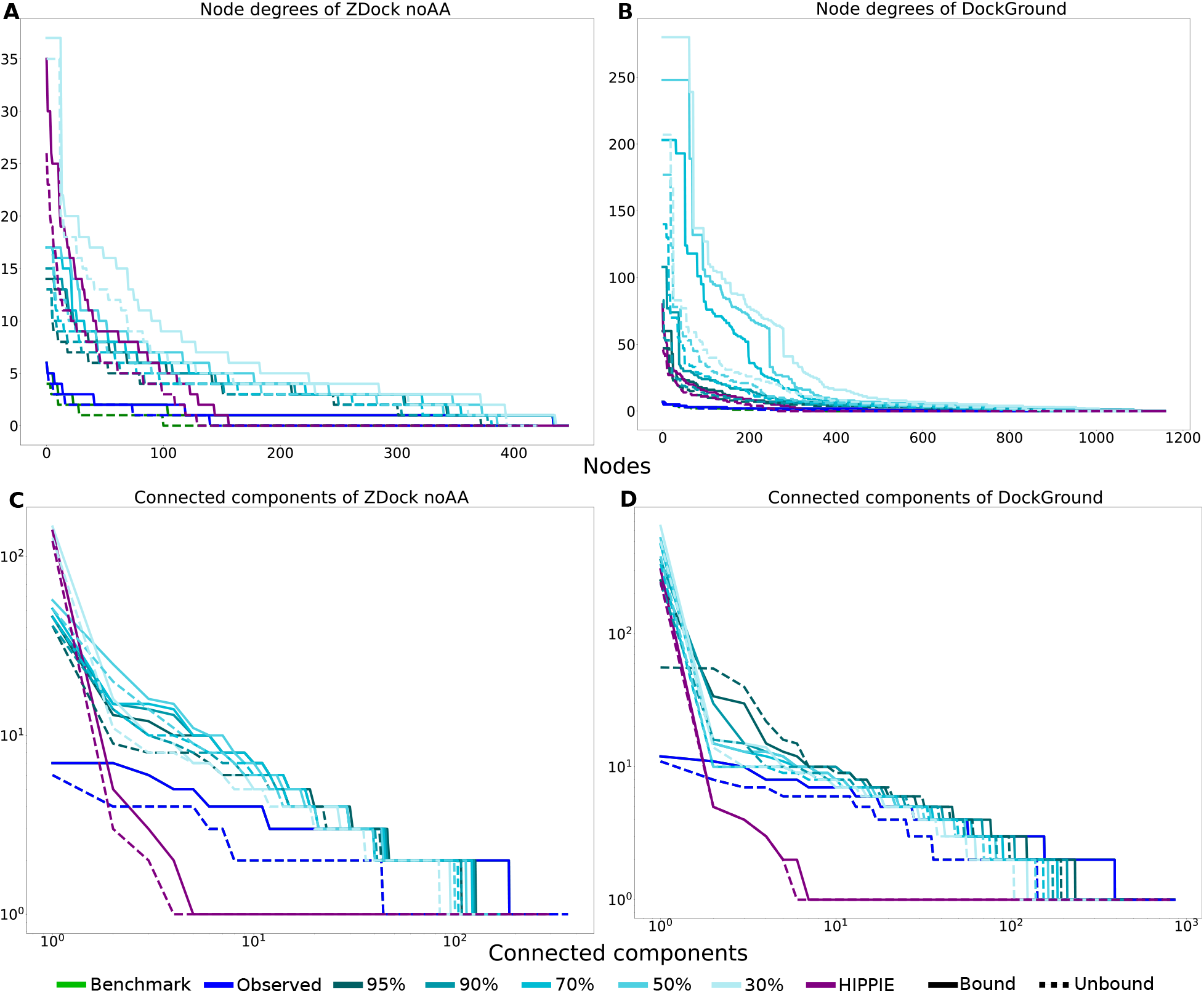
Node degrees and size of connected components in LEVELNET for ZDockv5 (antibody-antigens excluded) and Dockground databases. For each network, whose chains belong to ZDockv5 (**AC**) and Dockground (**BD**) databases, the node degrees (**AB**) and the size of connected components (**CD**) are sorted from the largest to the smallest. The x and y axis of the connected components are in logarithmic scale.

**Figure S3:**
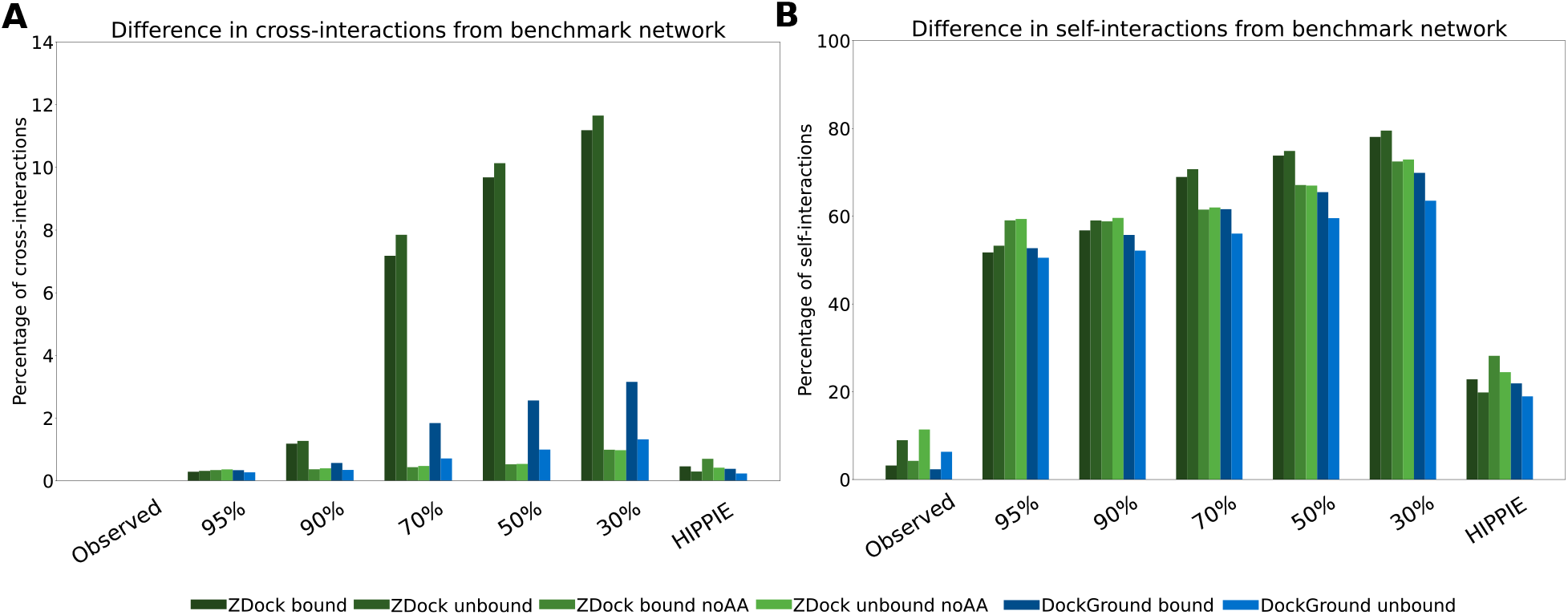
Differences of the interactions in each layer of LEVELNET against the benchmark network (measured in percentage of the maximum possible value). Two types of interactions are considered: (**A**) self- and (**B**) cross-interactions with the maximum possible values of *n* and 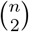 (fully connected graph), respectively.

**Figure S4:**
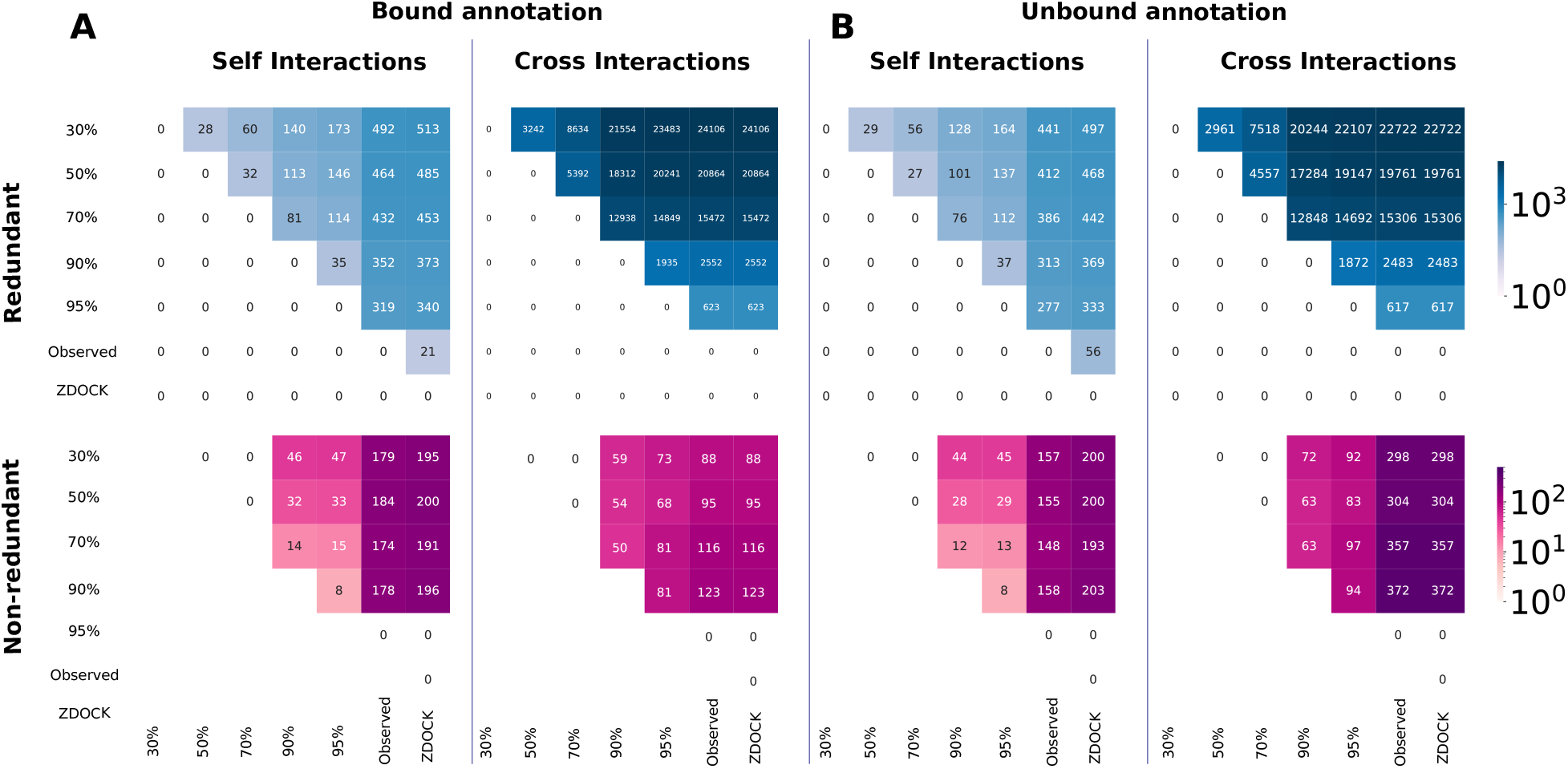
Pairwise differences in the number of interactions for layers of LEVELNET. In the case of redundancy reduction, each network has different set of nodes. In order to compare, for each cell, we took the network with smaller set of nodes (lower sequence identity) as the reference and compared it to the other network that is built based on the same reference nodes. There is no redundancy reduction for the physical contact and benchmark networks. Two types of interactions are considered: Self- and cross-interactions. **A**. ZDockv5 bound annotation. **B**. ZDockv5 unbound annotation.

**Figure S5:**
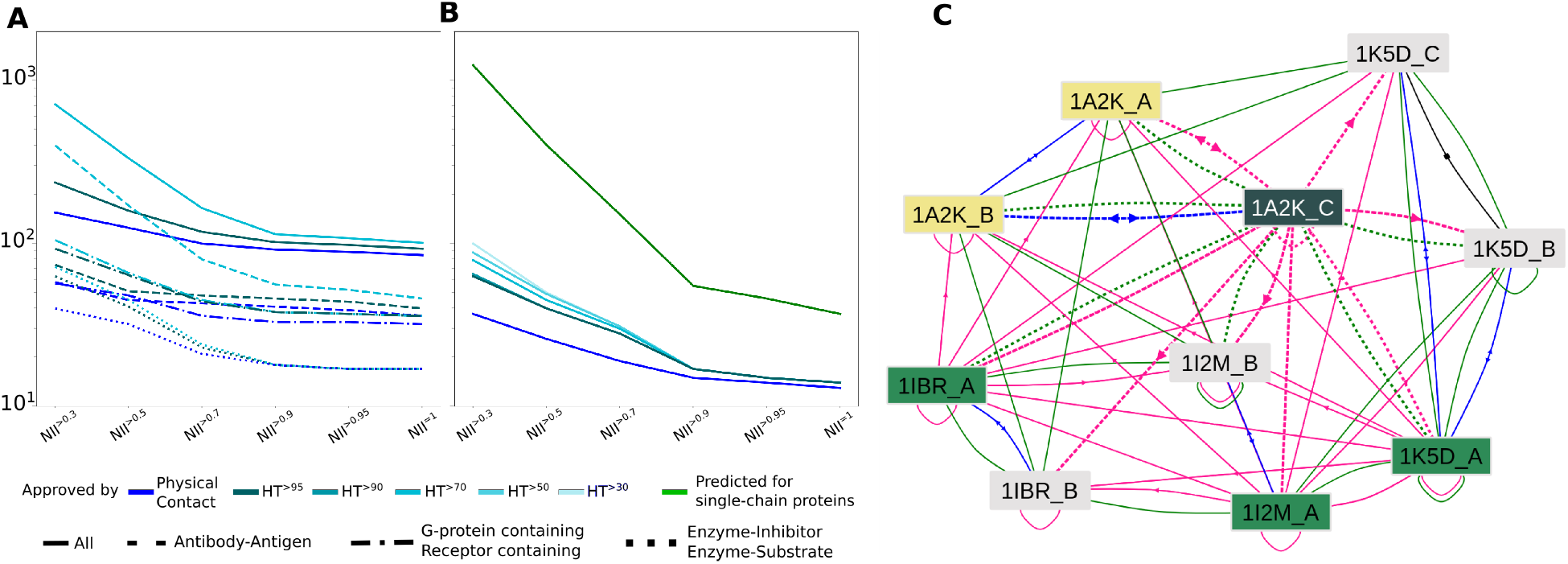
Assessments of PPI predictions. **AB**. Number of PPIs predicted for a set of 168 proteins ZDockv2 through complete cross-docking simulations [24] and supported by experimental structural data. The normalized Interaction Index (NII) is a confidence score assigned to each putative protein pair. Note that when a protein contains several chains, all paris of interacting chains are assigned a score of 1. We gave these scores as an input custom matrix to LEVELNET. The y axes are in logarithmic scale. **A**. The entire dataset and several functional classes are considered. **B**. Only the subset of 88 single-chain proteins is considered. **C**. Subnetwork for the GTP-binding nuclear protein RAN and its partners. The chains corresponding to RAN are 1A2K:C, 1IBR:A, 1K5D:A and 1I2M:A. Upon clicking on one of them, here 1A2K:C, LEVELNET interactive interface highlights the other ones, identified as homologs, in green. The other chains belonging to the same PDB complex are highlighted in yellow. The observed and homology-transferred (at >95% sequence identity) interactions are represented by the blue and pink edges, respectively. The green ones represent interactions predicted by cross-docking (taken from [22]. The interactions involving the selected chain (1A2K:C) are highlighted with dotted lines.

**Figure S6:**
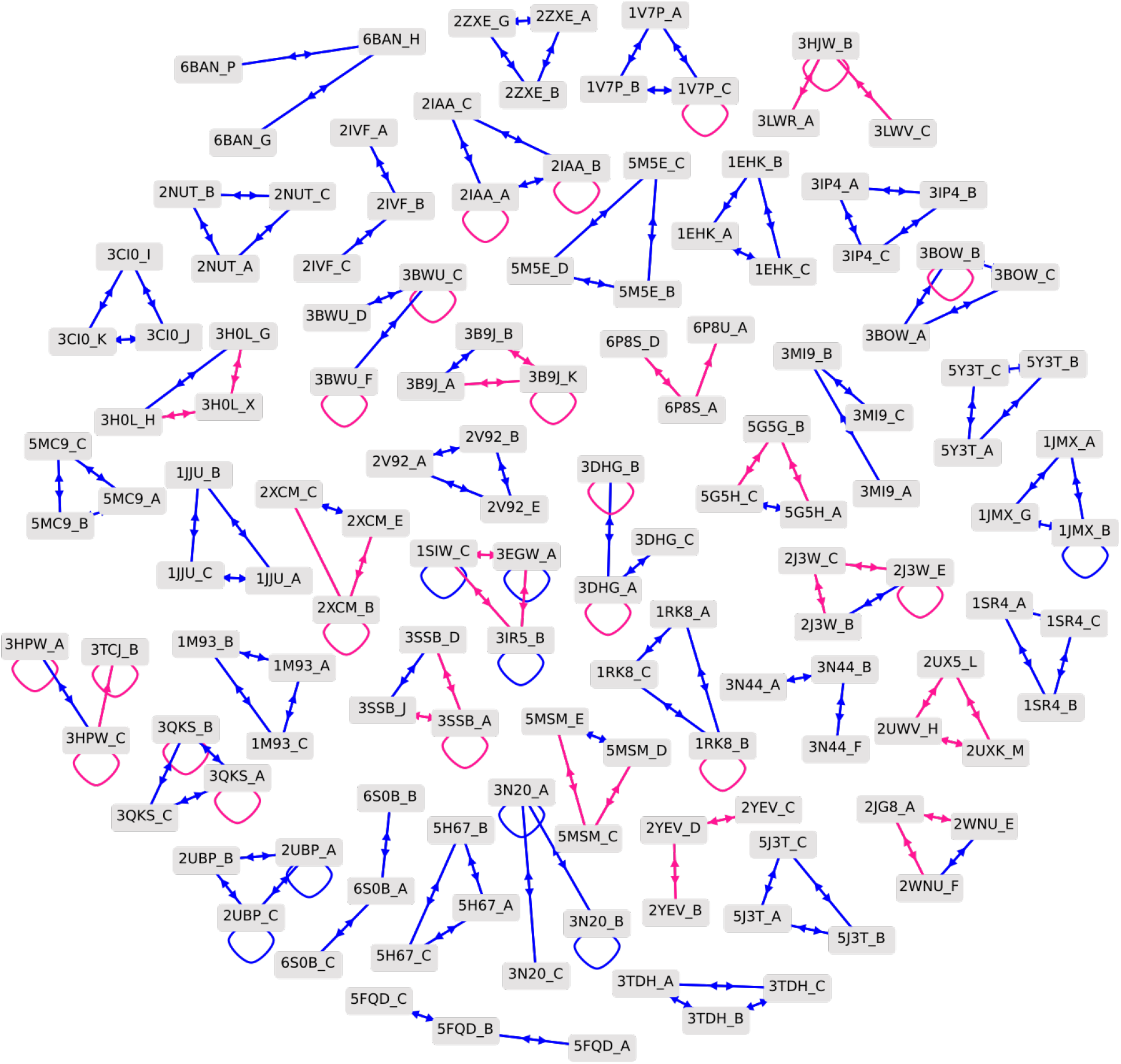
The PPI network of the new benchmark with 45 connected components of size 3 at sequence identity above 70%.

**Figure S7:**
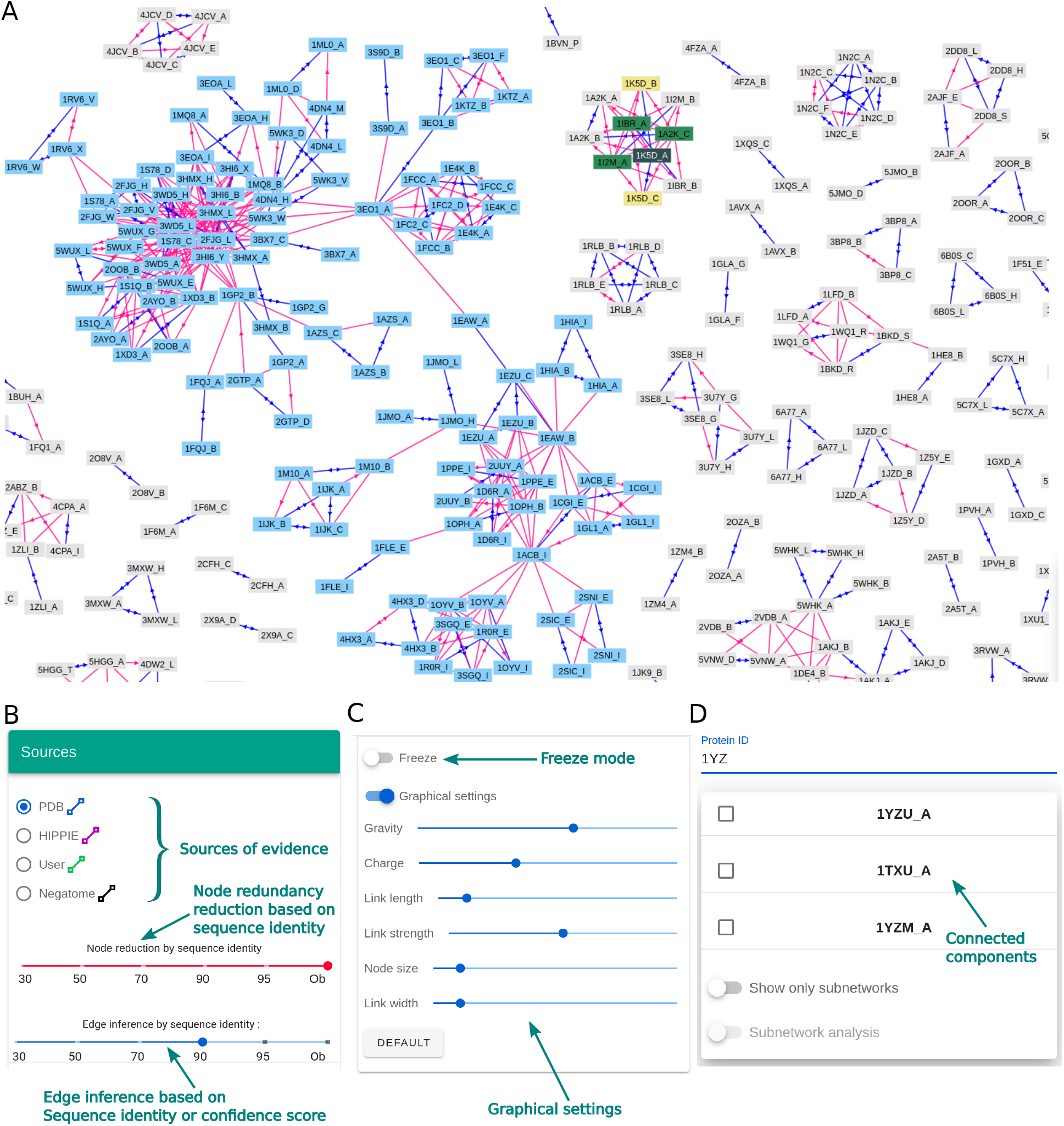
A snapshot of LEVELNET’s web-interface. **A**. Observed and inferred interactions with sequence identity above 95% of unbound single-chain proteins of the ZDockv5. Blue edges represent the existence of observed physical contacts between nodes. Pink edges are inferred by homology propagation from observed evidence. An arrow on an edge means that the partner in the destination is in a bound conformation. **AD**. The users can search and select a subset of homologous chains or connected components. **B**. The users can explore the network grids of different sources by modulating the node sequence identity and edge sequence identity (confidence score). **C**. Parameters to control the visualisation of the network.

